# Now What Sequence? Pre-trained Ensembles for Bayesian Optimization of Protein Sequences

**DOI:** 10.1101/2022.08.05.502972

**Authors:** Ziyue Yang, Katarina A. Milas, Andrew D. White

## Abstract

Pre-trained models have been transformative in natural language, computer vision, and now protein sequences by enabling accuracy with few training examples. We show how to use pre-trained sequence models in Bayesian optimization to design new protein sequences with minimal labels (i.e., few experiments). Pre-trained models give good predictive accuracy at low data and Bayesian optimization guides the choice of which sequences to test. Pre-trained sequence models also remove the common requirement of having a list of possible experiments. Any sequence can be considered. We show significantly fewer labeled sequences are required for three sequence design tasks, including creating novel peptide inhibitors with AlphaFold. These de novo peptide inhibitors require only sequence information, no known protein-protein structures, and we can predict highly-efficient binders with less than 10 AlphaFold calculations.

## I. INTRODUCTION

Sequence design is the construction of novel protein or peptide sequences which, when synthesized, will have chosen functional properties. Pre-trained sequence models like ESM,^1,2^ UniRep,^3^ and ProtTrans^4^ have recently shown excellent performance on structure prediction tasks with minimal experimental data. Reducing experimental data needed for sequence design is especially useful when finding peptides that bind to specific proteins, where predicting binding requires solid-phase peptide synthesis followed by assay.^5^ However, these pretrained models are not able to guide experiments – only make predictions with few data. Bayesian optimization (BO) is becoming the standard approach for choosing which experiments to do, but is incompatible with pretrained sequence models because BO requires accurate uncertainty predictions.^6^ We show how to modify pretrained sequence models to enable BO by modifying the encoder and readouts of the models. This marries the accuracy of pre-trained models with the ability to guide experiments of Bayesian optimization.

BO is a “black-box” optimization technique suited to expensive functions.^7^ BO can optimize a function without access to its derivative or any other information (the “black-box”) and optimizes with minimal evaluations of the function. BO has been successful in sequence, formulation, and molecular design problems where the “expensive function” is doing an experiment.^8–11^ BO is a *Bayesian* method, so it requires construction of a probability distribution surrogate model that approximates the black-box function (i.e., the experiment). BO works by choosing experimental data points in a way that balances **exploring** sequence space to improve accuracy of the surrogate model and **exploiting** the surrogate model to maximize the black-box function. The surrogate model is nearly always a Gaussian process regression model, ^6^ which is not as empirically expressive as neural networks^12^ and has limited ability to be pre-trained. We show here how to use pre-trained sequence models instead of Gaussian process regression. The advantage is that pre-trained sequence models are accurate with very few experiments and work well in the high-dimensional space of sequence design.

Current pre-trained models cannot be used as surrogate models in BO for two reasons. First, BO requires a gradient of the surrogate model with respect to its input, but sequence models have discrete integers as input. We use probabilistic reparameterization proposed in Linder and Seelig ^[13]^, which is similar to the Gumbel softmax-trick^14,15^ to enable gradient ascent of a pretrained sequence model. The second reason is that pre-trained models do not have uncertainties to compute probabilities in the BO algorithm. The classical solution to this problem is Bayesian neural networks,^16^ but Izmailov *et al.* ^[17]^ recently showed that deep ensembles^18^ are both competitive with integrating Bayesian neural network posteriors and more robust to noise in training data. We hypothesize that these two properties make deep ensembles a good approach in the low data regime targeted here.

Thus, our sequence design method is to modify pretrained sequence models by replacing the discrete input with a categorical distribution and deep ensembling to allow BO sequence design. In contrast to previous work,^19^ our method does not require a pool of known sequences and we can use any existing pre-trained sequence model without additional training. To test this method, we require the ability to label arbitrary new sequences (e.g., like you would in an experiment). We use here three tasks that mimic experiments to test our method: (1) designing a peptide that is hemolytic as evaluated by an RNN that is treated as a black-box;^20^ (2) matching an unknown sequence by only receiving BLOSUM-scored distance;^21^ and (3) designing a peptide that binds to a target protein as evaluated via an AlphaFold multimer calculation.^22,23^

### A. Related Work

Sequence design (also known as protein engineering) is a broad category of tasks like enzyme design, peptide drug design, design of self-assembled structures, and de novo protein design. There are multiple approaches to designing new sequences that range from purely experimental to purely computational. A commonly used experimental method is directed evolution,^24^ a process of optimizing fitness by stochastically mutating a wild-type sequence. Directed evolution typically requires a high-throughput assay that can select sequences. Directed evolution can be combined with computational approach.^25–27^ Cheng *et al.*^[28]^ proposed an efficient, experimental design-oriented closed-loop optimization framework for protein directed evolution, which employs a combination of novel low-dimensional protein encoding strategy and Bayesian optimization enhanced with search space prescreening via outlier detection. Harteveld *et al.*^[29]^ proposed a framework based to automatically assemble structural templates with native-like features. Our approach is differentiated from directed evolution because (1) our approach requires only testing a few dozen sequences and (2) the assay does not need to involve screening.

Machine learning for sequence design is beneficial especially where assays are expensive or slow enough to outweigh the cost and time of sequencing and synthesis.^30,31^ A recent review of *adaptive* machine learning approaches for sequence design can be found in Hie and Yang ^[32]^. Biswas *et al.* ^[33]^ used machine learning to guide the sequence searching by learning a latent representation from a small number of mutants. Bayesian optimization has been used for sequence design previously (without pretraining) in Hughes *et al.* ^[11]^ and Greenhalgh *et al.* ^[34]^. Khan *et al.*^[35]^ recently showed how to approach sequence design as a combinatarial Bayesian optimization problem with feasible trust regions for antibody design. Das *et al.*^[36]^ reported an efficient computational method for the generation of antimicrobials with desired attributes by leveraging guidance from classifiers trained on an informative latent space of molecules modeled using a deep generative autoencoder. Castro *et al.* ^[37]^ proposed a protein sequence design workflow by first training a deep Transformer-based autoencoder (ReLSO) to jointly generate protein sequences as well as predict fitness, following by fitness optimization over latent space and high fitness latent space sampling. Our work is similar to the previous Bayesian optimization approaches because it is iterative and calibrated, but different than approaches that are non-adaptive (train once on data).

Deep generative sequence models can be categorized into variational autoencoders (VAEs), generative adversarial networks (GANs) and language models, including RNNs and attention models. These are widely used to perform peptide generative with desired properies.^38–42^ Linder et al. ^[43]^ developed a generative method which can explicitly control sequence diversity during training by penalizing any two generated sequences on the basis of similarity. Ferruz *et al.*^[44]^ developed ProtGPT2, a Transformer-based generative model trained on UniRef50 which can generate de novo protein sequences following the principles of natural ones. Anand and Achim^[45]^ introduced a fully data-driven denoising diffusion probabilistic model for protein structure, sequence, and rotamers that is able to generate highly realistic proteins across the full range of domains in the Protein Data Bank by using equivariant transformers. These generative models can propose new sequences, but not inside an iterative BO algorithm like proposed here.

Pre-trained models for sequence design are increasingly common. Alley *et al.* ^[3]^ used an mLSTM model to learn statistical representations of proteins from 24 million UniRef50^46^ sequences. Rives *et al.*^[1]^ obtained sequence representations containing information about biological properties by using unsupervised learning to train a transformer language model on 86 billion amino acids across 250 million protein sequences spanning evolutionary diversity. Detlefsen *et al.*^[47]^ performed analysis on protein representations and demonstrated that pre-training representations can yield improved performance as well as significantly improve interpretability and let the models reveal biological information. Wang *et al.*^[48]^ reconstructed single-sequence 3D protein structure by feeding the pre-trained sequence embedding into a multi-scale network which is able to predict the interresidue 2D geometry. Transformer models equipped with self-attention mechanisms have shown to be particularly well-suited to capture dependency among sequence elements while being capable to scale vast amounts of model parameters.^49^ Kaplan *et al.*^[50]^, Hoffmann *et al.*^[51]^ indicate pre-trained sequence models could be compute-optimally trained by balancing compute, observations, and model parameters. Ruffolo *et al.* ^[52]^ used antibody affinity maturation with language models and weakly supervised learning. Russ *et al.*^[53]^ developed another sequence model for designing chorismate mutase enzymes. These promising results led to our choice of using pretrained model.

Bayesian neural networks have been proposed as an improvement to neural networks that account for uncertainty in predictions.^16^ The posteriors can be computed directly, with significant computational effort, via Hamiltonian Markov chain Monte Carlo (HMC).^54^ There are frequent efforts to reduce the cost with approximations, such as Maddox *et al.* ^[55]^ and specifically for chemical systems in Soleimany *et al.* ^[56]^. Deep ensembles^18^ are a common baseline that seems to be in practice as good as HMC while being more robust.^17^ Recent work has also proposed other ways of approximating uncertainty in transformers, the most common architecture for pretrained sequence models.^57^

## II. THEORY

The labeling of a sequence *x* with its property *y* is done with an expensive black-box function *f*(*x*). *f*(*x*) could require synthesis prior to an experiment, a molecular dynamics calculation, or an expensive calculation. Our goal is to find *x\** that maximizes *f*(*x*\*) through BO starting with zero labeled data points. We indicate iterations through the BO algorithm with index *n*(*x_n_)*

We construct a pre-trained sequence model that embeds a sequence *x* ∈ {0, 1}^*L*×*A*^ into a continuous vector space 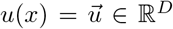, where *L* is the sequence length, *A* is the number of possible tokens in the sequence (alphabet), and *D* is the dimension of the sequence representation. We use UniRep for *u*(*x*).^3^ Unirep is a long short-term memory (LSTM) model trained to perform next amino acid prediction, as implemented in JAX.^58,59^ Properties are predicted from 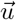 with a multi-layer perceptron (MLP) 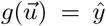. To enable uncertainty predictions, we use deep ensembles.^18^ Deep ensembles predict a normal distribution parameterized from *M* models 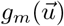. Finally, to enable optimization of the input sequence x, during BO we sample the sequence from a categorical distribution characterized by trainable logits, similar to the Gumbel-Softmax Trick.^14,15^ These individual components are described in more details below and visually in Figure 1.

**FIG. 1.**
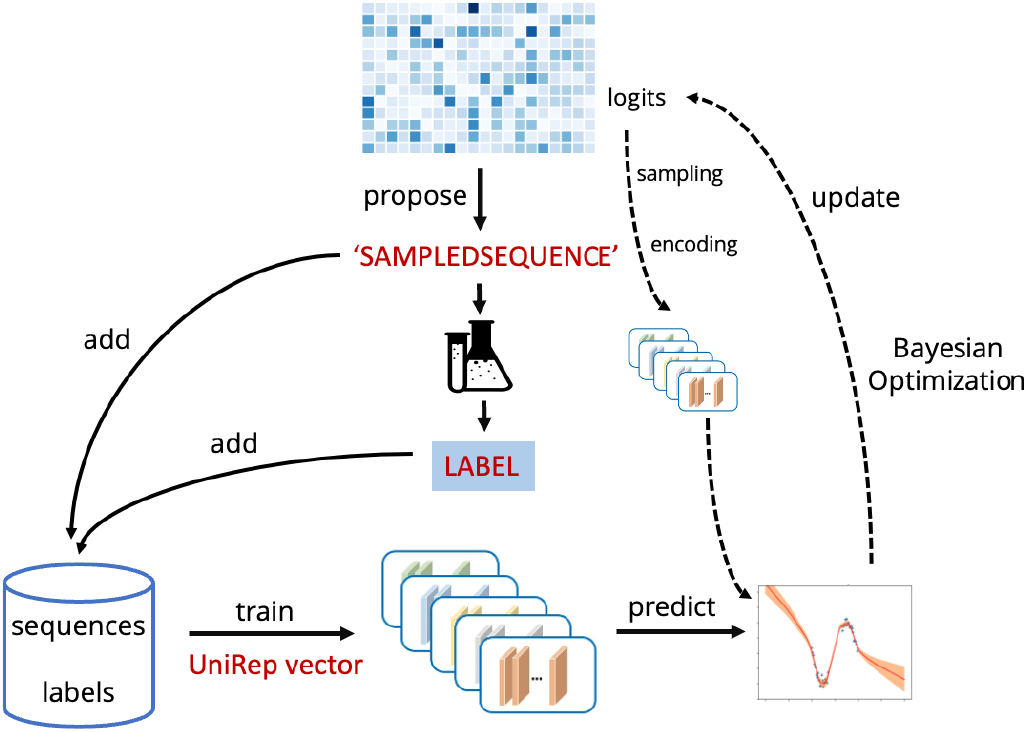
An overview of the model and Bayesian optimization process. Sequences are defined by logits during BO acquisition function maximization and then labeled. The complete set of sequences and labels is then re-used to train the deep ensembled MLP. Finally, the MLP is used for the next round of BO.

### A. Model

UniRep^3^ is an LSTM model^60^ trained on Uniref50^61^ to perform next amino acid prediction by minimizing crossentropy loss. By conducting this semi-supervised classification, the model learns how to internally represent protein sequences. Given a sequence of any length, UniRep returns a single fixed-length vector representation. We denote UniRep as 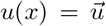, where *u*(·) is the UniRep model, which takes sequence *x* as input and outputs the representation vector 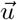. The dimension of 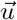 is *N* = 1900 for UniRep. See Alley *et al.* ^[3]^ for complete details. We use a JAX implementation of UniRep.^58,59^

Uncertainty in predicted labels can be split into an epistemic uncertainty (EU) and aleatoric uncertainty (AU).

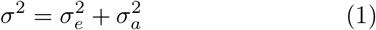

where 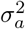; is AU and 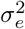 is EU. AU is also known as data uncertainty or statistical uncertainty. It is unavoidable noise from *f*(*x*). For example, data collected from a laboratory will have uncertainty. EU is also known as model uncertainty or system uncertainty. It arises from the lack of knowledge of the system in respect to quantities and processes within the system. It can be reduced through larger or better models. High EU arises in regions where there are few or no observations for training. The output from each single MLP *g_m_*(*x*) is two numbers that characterize a normal distribution 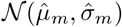. We can use these to provide estimates of the AU and EU:^18^

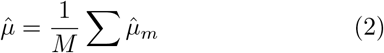

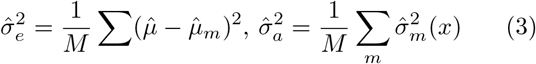

Where *M* is the ensemble size, *m* is the index of the MLP.

Likelihood describes the joint probability of the observed data as a function of the parameters of the MLPs. Negative log-likelihood minimization is a proxy problem to the problem of maximum likelihood estimation. We assume that the labels *y* follow a normal distribution of *P*[*f*(*x*) = *y*] characterized by *μ*(*x*), *σ*(*x*). The negative log-likelihood of model parameters is the probability of observing the label *y* given sequence *x*:^18^

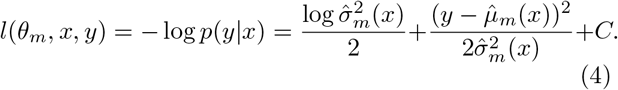

As proposed in Lakshminarayanan *et al.* ^[18]^, we also do adversarial training. After one step with the loss in Equation 4, we perform another step to smooth the model predictions:

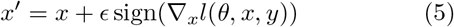

where *ϵ* is a hyperparameter that controls the strength.

### B. Bayesian Optimization

Bayesian optimization is a gradient-free global function optimization method which is constructed for expensive-to-evaluate functions.^6^ The goal of Bayesian optimization is to both explore and exploit existing knowledge as expressed in an acquisition function. Namely, our next sequence to test is computed from

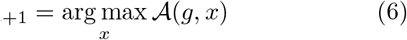

where 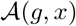 is our acquisition function that uses the model (*g*) and sequence (*x*). We use the simplest acquisition function for balancing exploration and exploitation called upper-confidence bound (UCB):

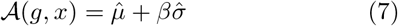

where *β* is a hyperparameter that balances exploration and exploitation that can be scheduled to increasingly exploit over the BO algorithm. UCB is not robust to label noise, and recent work has proposed different acquisition functions robust to label noise.^62–64^ To alleviate label noise, we only use EU 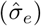 in Equation 7. We chose UCB due to its simplicity and robust performance across hyperparameter choices.

BO requires computing a gradient of the surrogate model – UniRep – with respect to input x to maximize the acquisition function.^65^ This is problematic, since *x* is not continuous. We can redefine *x* as being a random categorical distribution parameterized by continuous logits *l* ∈ ℝ^*L*×*A*^. Then when a sequence is needed, *x_i_* is computed from a random drawn categorical at the ith position (where *i* indexes positions in sequence and *j* indexes amino acid/word):

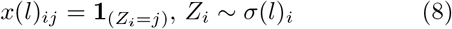

where *σ*(*l_i_*) is the categorical distribution parameterized by *l_j_* This reparameterization makes the gradient accessible, but we must propagate a gradient through sampling from the distribution. We use a straight-through approximation

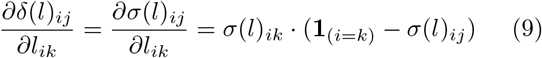

However, the logits can drift close to zero so that the drawn sequences are highly variable or the logits can grow in magnitude and gradient updates no longer actually change the sequence. The standard solution to this is the Gumbel-Softmax Trick.^15^ We used a slightly different approach introduced by Linder and Seelig ^[13]^, which is to simply add a trainable layer normalization that can trainably affect the mean and variance of logits. ^66^ Recently, Daulton *et al.* ^[67]^ showed that BO with probabilistic reparameterization, like Equation 8, will converge to the true maximum of the acquisition function. Although, convergence of the *L* × *A* logit matrix is still a difficult high-dimensional optimization.

We considered using the continuous latent space as well. In Figure S1 we use the UniRep decoder to avoid optimizing the sequence directly. We optimized 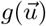 by working with 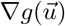. We found that the decoder 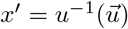 gave a sequences back that were inconsistent enough with the forward label *g*(*u*(*x*′)) and prevented optimization past a certain point. Figure S1 shows optimization of latent space has continuous improvement, but after decoding to an actual sequence and evaluating 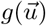 there is a plateau. This may be unique to UniRep or because or be task specific because other recent work has successfully used latent spaces for BayesOpt^68,69^ and work on improving extrapolation from latent spaces.^70^ Nevertheless, avoiding latent space negates this potential agreement problem between 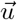 and *x*.

Variable sequence lengths were incorporated by maximizing the acquisition function over lengths *L* – 1, *L*, and *L* + 1 at each BO iteration and the length L was replaced by the best for the next iteration.

## III. METHODS

The deep ensembled MLPs are sensitive to mode collapse, where all *g_m_*(*x*) models converge to the same parameters by overfitting the training data.^71^ This is common in our setting of only a few data points. We use three strategies to mitigate this. First, we use relatively few parameters in the MLPs to frustrate the loss landscape and make it sensitive to initial parameters. Second, we resample data so that the training data seen by each MLP is different. Lastly, we employ standard techniques to reduce overfitting like dropout and weight decay. There are more systematic approaches that could be used,^72^ but we found these simple strategies give good calibrated uncertainties across the three tasks. Adversarial training was found to have minimal improvement, even when adding significant artificial label noise (Figure S3). All hyperparameters, including architecture and training parameters are given in Table I.

**TABLE I.** Hyperparameters for work presented here. All were tuned on finding unknown target peptide sequence task.

| Hyperparameters | Choice |
| --- | --- |
| MLP shape | 128, 32, 2 |
| Activation | Swish<sup>73,74</sup> |
| Ensemble number | 5 |
| training epochs | 100 |
| training batch size | 8 |
| training learning rate | 1e-4 |
| training resampled classes | 10 |
| training weight decay (Adam) | 0.1 |
| training $\epsilon$ | 0.001 |
| dropout rate | 0.2 |
| BO batch size | 16 |
| BO iterations | 200 |
| BO $\beta$ | 2.0 |

While training, we found resampling data to account for non-uniformity in label distribution is important – especially because BO targets high values that are rare. So while training the model, we resampled training data according to their labels by binning training data into 10 classes and sampling with replacement to get equal frequency. As discussed above, we repeated this process for each MLP (*g_m_*(*x*)), so that they saw different training data frequencies.

Maximizing Equation 7 for BO is a non-convex problem^75^ and so we started from 16 initial logit distributions and performed 200 gradient ascent steps with Adam^76^ (Figure S4). This was repeated for each considered length - one higher and one lower than current length. Daulton *et al.*^[67]^ recommended L-BFGS, although we found Adam works for our tasks as shown in Figure S4 (effect of increasing number of starting points/iterations).

## IV. RESULTS

We tested our algorithm on three different tasks: designing a hemolytic peptide, finding an unknown target sequence, and designing a peptide that binds to Ras GTPase.^77^ In the first two tasks, we also compared with ablations by removing the pre-trained model with one-hot encoding and/or using greedy optimization instead of BO.

### A. Hemolytic peptide

The first task is to design a hemolytic peptide. Hemolytic peptides are small 2-80 peptides that lyse red blood cells. Predicting hemolysis is important when assessing if a peptide will make a viable therapeutic. In place of an experiment, we use a previously trained model as an stand-in for *f* (*x*). This model is a bidirerctional LSTM^20,60^ that predicts of a sequence will be a hemolytic peptide. It is trained on data from Pirtskhalava *et al.*^[78]^. This task has variable sequence length and is initialized to a random peptide with random lengths from 10–20 uniformly sampled. Figure 2 shows the results of our algorithm averaged (mean) over 50 independent runs. Figure 3 shows corresponding distribution of sequence lengths over the course of the algorithm. It can find a likely hemolytic peptide within 5 iterations and nearly match the most hemolytic predicted peptide from the 9,316 peptides analyzed in Ansari and White^[20]^ after 20 iterations.

**FIG. 2.**
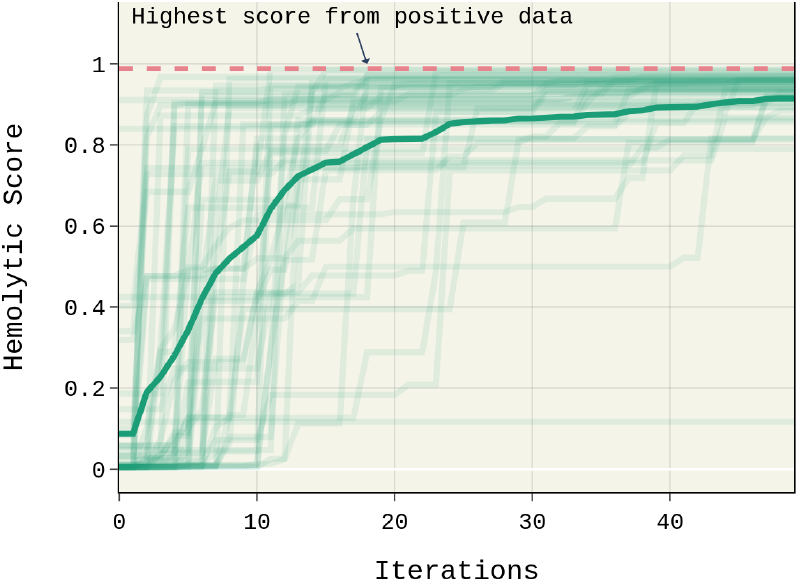
Current best sequence during iterative optimization averaged across 50 runs. Different traces show individual runs. The solid line show the result averaged(mean) over 50 individual runs. Sequence length is allowed to change here. Iterations is same as number of sequences observed.

**FIG. 3.**
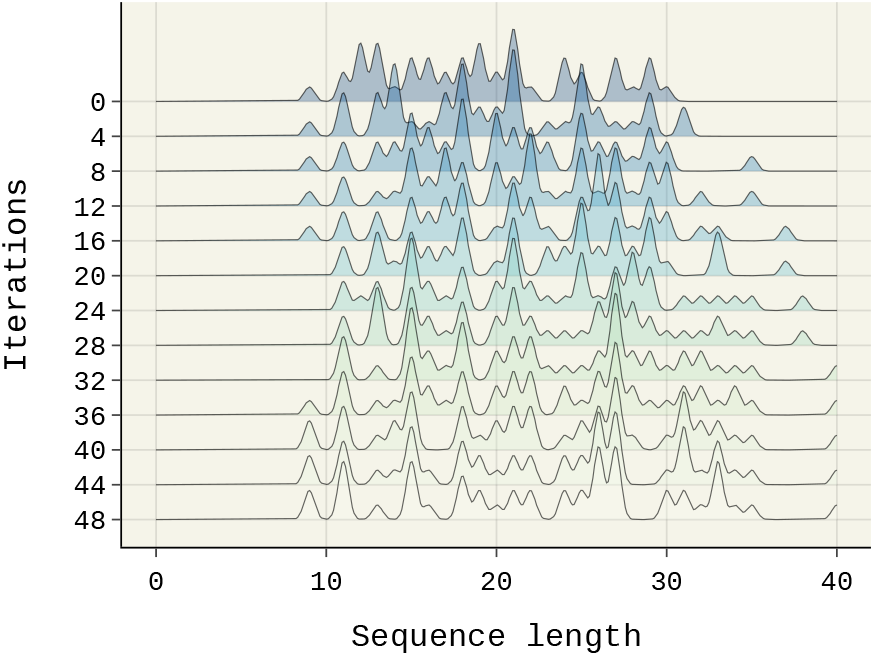
Kernel density estimates of sequence length from Figure 2 algorithm as a function of iteration number. Initial sequence length is randomly sampled from a uniform distribution of [10, 30]

Figure 4 compares against two ablations, showing the gain from adding pre-training and BO where the sequence length is not allowed to change due to the use of one-hot encoding. We find that indeed both components of our algorithm help in this task, and pre-training is better at all iterations.

**FIG. 4.**
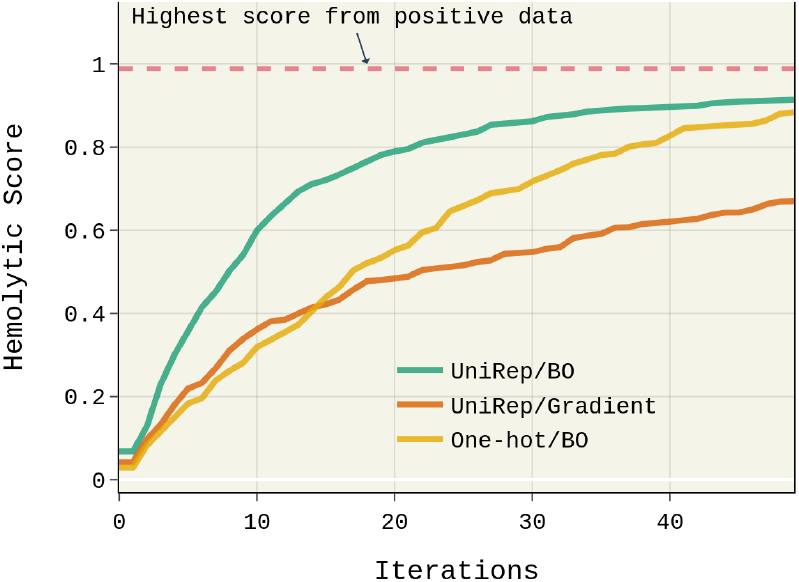
Current best sequence during iterative optimization averaged across 50 runs. The different lines are the algorithm presented here (UniRep/BO) and ablations. Sequence length is fixed to thirteen to compare with one-hot encoding. Iterations is same as number of sequences observed.

### B. Unknown Target Matching

In the second task, the sequence length is fixed at thirteen residues. *f* (*x*) is the similarity score between *x* and an unknown target. Similarity is measured by BLOSUM62 matrix,^21^ which gives disagreement weighted by evolutionary data. This makes it so chemically similar side-chains disagreeing is less important. This task is extremely specific, so we would expect pre-training to be minimally effective or even prevent learning. This task represents a worst-case for pre-training.

Model hyperparameters in this work have been tuned for scores that approximately range from 0-10 and so we transformed the score according to:

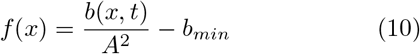

where *b* the BLOSUM62 score between *x* and the target *t* and *b_min_* is the minimum possible score. The calibration (correctness of uncertainty) of the MLP is shown in Figure S2.

Figure 5 shows the results of the proposed algorithm and ablations. This task is convex (no local maximums) and has no noise, so BO is similar to direct gradient ascent of the surrogate model predictions. One-hot is direct use of the sequence in the MLP without pretrained model. We expect this to be a better representation, since our task is highly specific, and indeed with enough data it eventually surpasses pre-training. Nevertheless, pre-training shows significant gains with fewer data points.

**FIG. 5.**
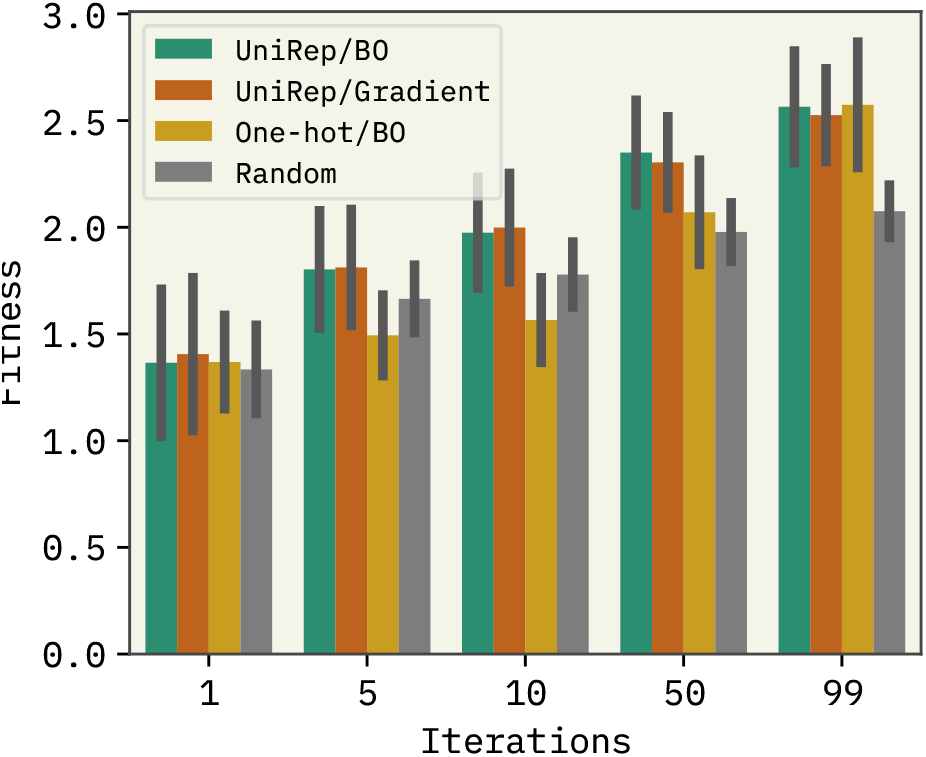
Current best sequence on unknown target matching task along with ablations. BO is similar to direct gradient ascent (greedy) of surrogate model because this task is convex and has no noise. Pre-training helps at low iterations, but is surpassed by one-hot at later iterations. This is expected on this highly-specific task. Error bars are 95% confidence intervals from bootstraping 50 repeated BO runs.

### C. AlphaFold2 protein-peptide binding

This task is to identify a candidate peptide that binds to a target protein as evaluated with AlphaFold2.^22^ The target protein is encoded by oncogene KRas G12C associated with development of cancer.^79^ Margarit *et al.*^[77]^ showed that activation of Ras GTPase is catalyzed by nucleotide exchange factor Son of Sevenless (SoS). As a result, an effective SoS inhibitor that would bind to the receptor-binding domain of the oncogenic system and preventing Ras overexpression is a therapeutic target. Konstantinopoulos *et al.*^[80]^ summarized findings that two important regions (based on conformational changes) are residue indices 24 to 40 and 56 to 75.

We specifically used AlphaFold2-Multimer,^23^ since this task is to predict simultaneously the Ras GTPase and bound peptide. Isak Johansson-Akhe^[82]^ showed that AlphaFold2-Multimer has accuracy similar or better than other docking programs in predicting peptide-protein complexes from scores on known complexes. Using this result, Chang and Perez^[83]^ showed a novel application of AF2-Multimer for competitive binding of different peptides to the same receptor. Following their work, we correlate good binding with the “confidence” score called predicted local distance difference test (pLDDT) output by AlphaFold2 and measuring distance to the binding site of SoS:

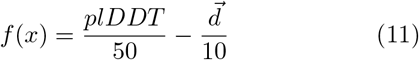

where 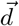 is the mean distance of the peptide to the binding site evaluated at the closest backbone atom per residue pair between peptide and Ras GTPase. The linear combination of 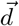 and pLDDT enables simultaneous minimization of mean distance and maximization of pLDDT, and keeps the score magnitude between 0 to 10. We use this for simplicity, although there are more principled methods for multi-objective BO.^84^ There are also other approaches to finding peptide binders that do not involve BO.^85,86^

Figure 6 shows the BO of Equation 11 averaged across 5 runs with variable sequence length. We compared against results from Anupam Patgiri^[81]^, which experimentally found the peptide FEGIYRLELLKAEEAN to bind well to Ras GTPase. They experimentally verified the peptide binds to the two important sites (I 25-40, II 56-75) and we found our binding protocol indeed shows binding near site I. When restricting the length to be the same as the Patigiri sequence, the sequence that maximized Equation11 is DANKEQMAQARQRAKQ. Its average distance from site I is 3.4Å and plDDT is 68. The Patgiri sequence is 5.4Å and 37 respectively. It is predicted to be a tighter binder than the Patgiri sequence. Of course, this requires experimental validation.

**FIG. 6.**
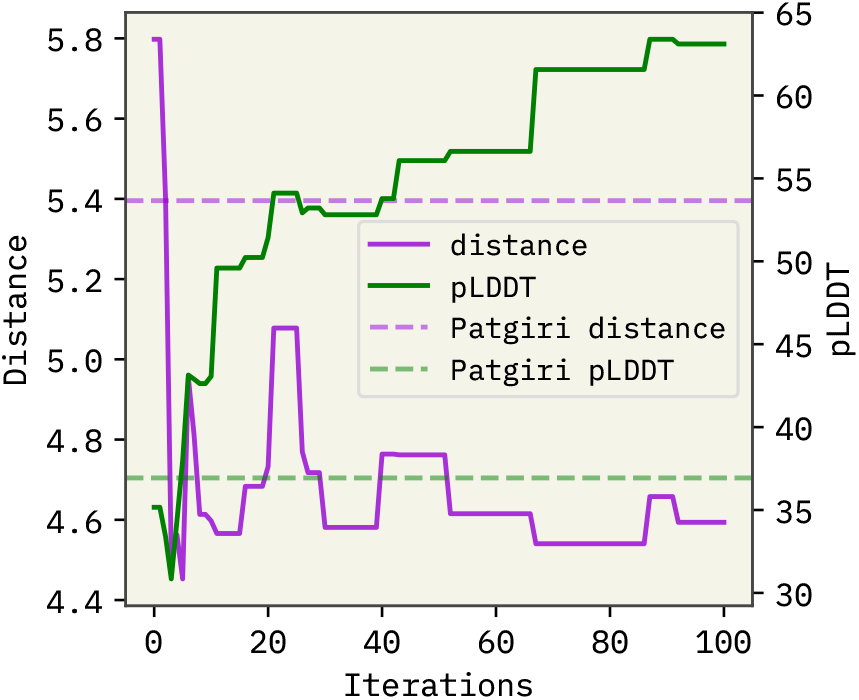
Average of 5 runs of algorithm on AF2 task. The dashed line shows average distance and pLDDT of the known binder to Ras GTPase. BO optimization finds a better peptide within a few iterations.

Figure 7 shows a comparison of the output from a BO run vs the Patgiri binder. It is clear that the peptide binder is closer - agreeing with the score. The sequence features strong *α*-helix formers and puts alanines near the binding site I, which creates a hydrophobic packed region. The combination of this hydrophobic core and strong helix gives it an exceptionally-high pLDDT score. It is not known if this will translate to a better affinity in an experiment.

**FIG. 7.**
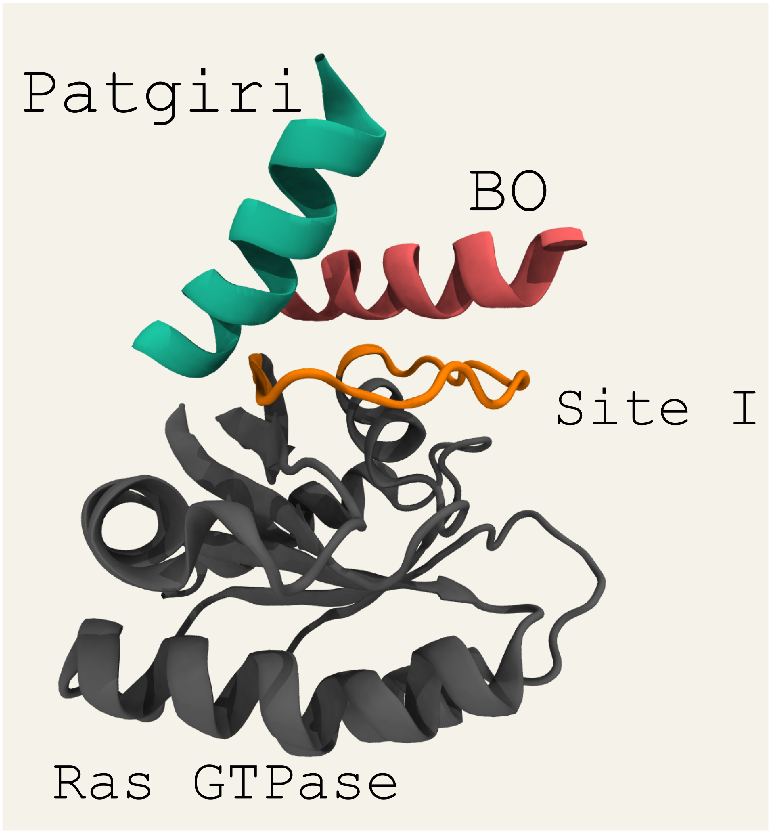
Comparison of BO optimal peptide binder and the AlphaFold2-Multimer prediction of the Anupam Patgiri^[81]^ (Patgiri) peptide binder. The BO sequence is DANKEQ-MAQARQRAKQ.

## V. CONCLUSIONS

We have shown how to use pre-trained sequence models in a BO algorithm. Across three tasks, this process enables good optimization with only a few data points. Our strategy was deep ensembled MLPs to provide calibrated uncertainties and probability distributions over sequence space to enable end-to-end differentiation. We found the commonly proposed optimization in latent space followed by decoding does not work well with few examples. We proposed using this in conjunction with AlphaFold to design de novo peptide binder from sequence alone.

## ACKNOWLEDGMENTS

Research reported in this work was supported by the National Institute of General Medical Sciences of the National Institutes of Health under award number R35GM137966. We thank the Center for Integrated Research Computing (CIRC) at University of Rochester for providing computational resources and technical support.

## CODE AVAILABILITY

Code is available at https://github.com/ur-whitelab/wazy. Hemolytic training data is available at https://github.com/ur-whitelab/peptide-dashboard.

## Notes

https://github.com/ur-whitelab/wazy

### Competing Interest Statement

The authors have declared no competing interest.

### Summary of Updates

Changed peptide binding formula and updated correspondingly Figs 6-7. Fixed ordering of figures and added additional citations.

