## Supporting Information for "Now What Sequence? Pre-trained Ensembles for Bayesian Optimization of Protein Sequences"

(Dated: September 3, 2022)

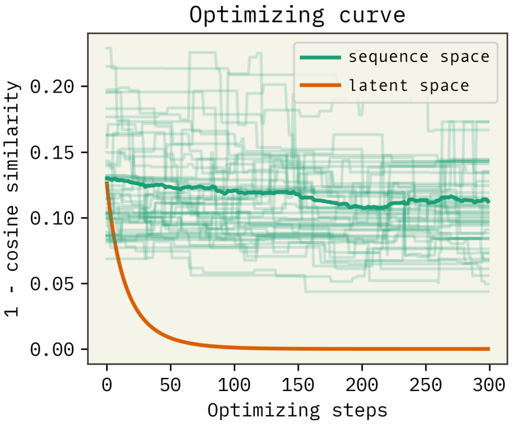


FIG. S1. Optimization of $1 - \vec{u} \cdot \vec{t}$ where $\vec{t}$ is a constant via working with UniRep latent space $u(x)$ directly. The organge line shows an optimal $\vec{u}$ is found quickly. The green line shows that recovering the sequence for the optimum via decoding $u^{-1}(\vec{u})$ gives a sequnce that, when encoded, is not actually improving.

---

*

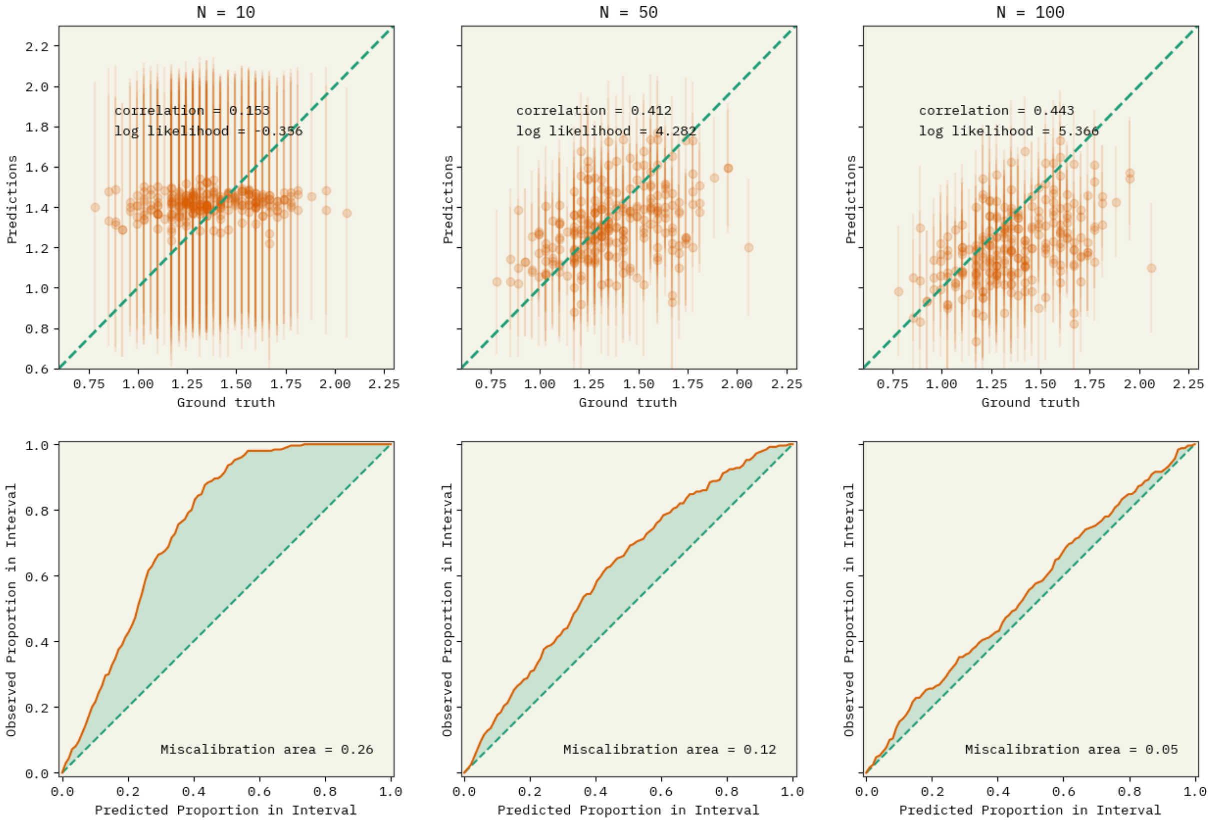


FIG. S2. Predictive model performance over amount of training data, 10, 50, 100 respectively from left to right show improvement in calibration and accuracy. The first row shows parity plots with uncertainty shown as error bar (95% confidence interval from $\hat{\sigma}(x)$). The second row shows fraction of labels that fall within predicted score.

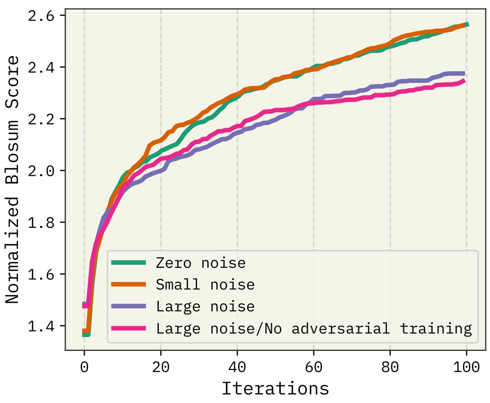


FIG. S3. Model performance on BLOSUM task (task 1) with increasing amounts of noise and without adversarial training. Adversarial training should make the model more robust to noise, but seemed to have little effect even with large noise.

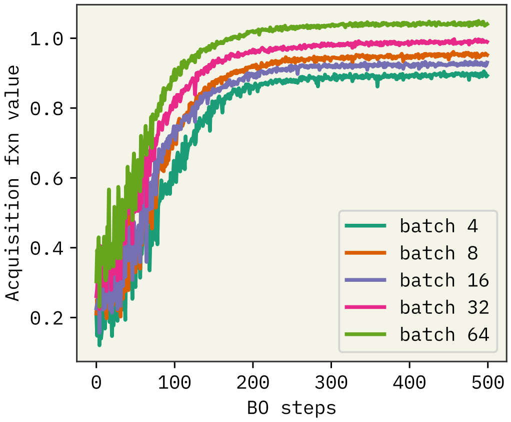


FIG. S4. Convergence of Acquisition function (UCB) as a function of stochastic gradient descent steps and batch size. This is from the hemolytic task. Based on these results, batch size 16 and 200 steps was chosen.

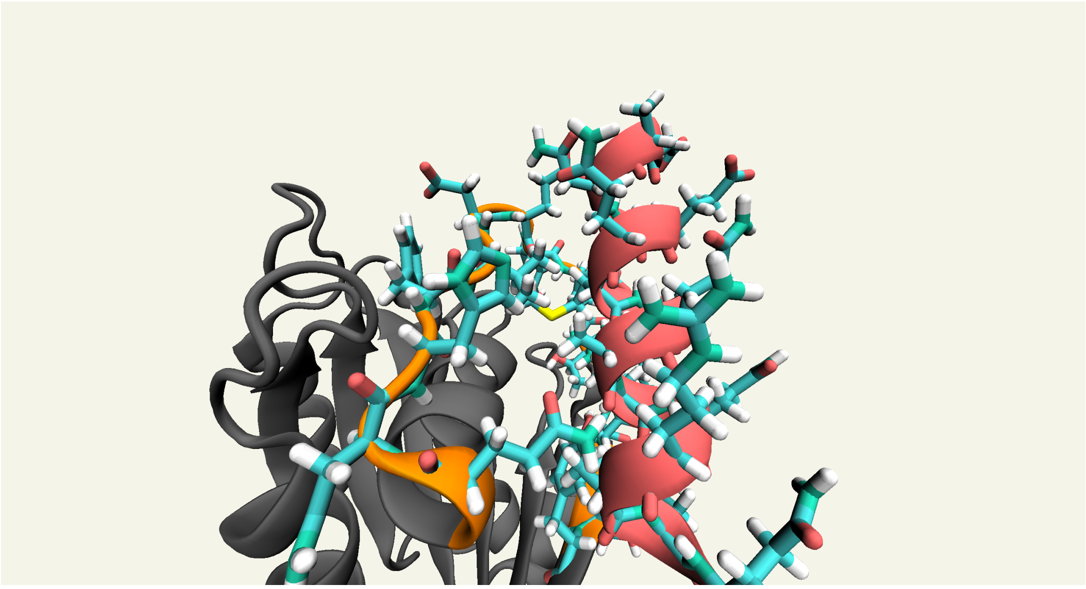

FIG. S5. Explicit side-chain atoms of Site I and optimized peptide binder DANKEQMAQARQRAKQ. Shows alanine fomrning hydrophobic area around Site I.
